# Decoding directional genetic dependencies through orthogonal CRISPR/Cas screens

**DOI:** 10.1101/120170

**Authors:** Michael Boettcher, Ruilin Tian, James Blau, Evan Markegard, David Wu, Anne Biton, Noah Zaitlen, Frank McCormick, Martin Kampmann, Michael T. McManus

## Abstract

Genetic interaction studies are a powerful approach to identify functional interactions between genes. This approach can reveal networks of regulatory hubs and connect uncharacterised genes to well-studied pathways. However, this approach has previously been limited to simple gene inactivation studies. Here, we present an orthogonal CRISPR/Cas-mediated genetic interaction approach that allows the systematic activation of one gene while simultaneously knocking out a second gene in the same cell. We have developed this concept into a quantitative and scalable combinatorial screening platform that allows the parallel interrogation of hundreds of thousands of genetic interactions. We demonstrate that the established platform works robustly to uncover genetic interactions in human cancer cells and to interpret the direction of the flow of genetic information.

## Introduction

For over a decade, RNA interference (RNAi) has been used to systematically assign function to genes by studying loss-of-function phenotypes^1^. More recently, it was shown that the bacterial pathogen-defence system CRISPR/Cas can be used to edit mammalian genomes^2^ and carry out a large variety of additional tasks including transcriptional repression and activation as well as epigenetic modifications^3^. The currently available CRISPR/Cas toolbox allows the highly parallel functional interrogation of every single gene in the human genome^4^. Nevertheless, dissecting the functional connectivity within complex cellular circuits is a major challenge in mammalian biology. While knockout and knockdown approaches above have proven highly successful to systematically attribute function to individual mammalian genes, they do not provide a deeper understanding of how these genes function together in complex genetic signalling networks. For this purpose, genetic interaction mapping has proven highly valuable to assign functional connectivity between cellular components.

Genetic interaction mapping traditionally analyses loss-of-function phenotypes of individual genes and their combinations to identify aggravating or alleviating genetic interactions^5^. Genetic interaction studies in model organisms including yeast^6-10^, C. elegans^11^, Drosophila^12^, and more recently human cells have established functional relationships between genes^13-18^. Although these studies have illuminated new connections between genes, they have not explicitly addressed the directionality of genetic interactions in hierarchical genetic networks in human cells. But, knowing the direction of genetic interactions can be critical for properly interpreting functional dependencies between genes, and offer rational approaches for therapeutic intervention. Yet, attempts to illuminate the direction of genetic interactions have been limited to model organisms^19-22^. In these studies, directionality could only be inferred in cases where either one of the interaction partners displays a no-loss-of-function phenotype, or both partners display opposing phenotypes^20^. However, the loss of each of two interacting genes generally has a phenotype by itself^19^ and in the case of activating interactions, which are frequently found in aberrantly activated signalling cascades in cancer cells, such as MAPK signalling^23^ - these phenotypes go in the same direction.

Overall, tumour genomes display a large variety of genetic and epigenetic changes, causing tumour initiation, progression and therapy resistance^24^. Owing to next-generation sequencing technology, the list of genes that are characterised to be either mutated or differentially expressed in different tumours is growing steadily^25^. The functional interpretation of such sequencing data, however, is challenging and further complicated by the fact that genes rarely ever act by themselves, but rather in complex genetic interaction networks, in which the function of one gene depends on the status of interacting genes^26^.

To reconstruct directional regulatory networks in human cells, we developed an orthogonal CRISPR/Cas system composed of two Cas9 enzymes from different species. This system allows the simultaneous and asymmetric activation of one gene and knockout of a second gene in the same cell. When compared to conventional symmetrical loss-of-function experiments in which the function of both interaction partners is lost, our orthogonal asymmetric platform allowed us to determine whether the activated gene functionally depends on, or can compensate for the loss of a knocked-out gene. Using this platform, we identified directional genetic interactions between genes whose activation or knockout altered the fitness of human chronic myeloid leukaemia (CML) cells. We demonstrate that the orthogonal screening approach can quantify loss- and gain-of-function phenotypes from the same cell, and that it is suitable to systematically identify genetic interactions between cancer relevant genes. Importantly, we managed to reconstruct a significant number of directional dependencies, connecting uncharacterised ‘dark matter’ genes to well-studied pathways.

## Results

### CRISPRa screen identifies cancer pathway genes

CML is a leukaemia characterised by a reciprocal translocation between chromosome 9 and 22. This translocation creates the BCR-ABL fusion oncogene, a constitutively active tyrosine kinase oncogene that causes myeloid precursor cells to divide in an uncontrolled fashion^27^. Application of BCR-ABL tyrosine kinase inhibitors (e.g. imatinib) have revolutionised treatment for this cancer, and decades of study have yielded fundamental information on the genes critical for BCR-ABL dependent signalling. We thus chose CML to benchmark our novel directional genetic interaction platform, using the K562 CML cell line to systematically identify genes that function as negative or positive regulators of cancer cell fitness. We found that imatinib can tune the viability of K562 cells over a broad range of concentrations (10 - 1,000 nM), and that CRISPR mediated activation (CRISPRa) of the imatinib efflux transporter ABCB1 resulted in an approximately 2-fold increased IC_50_ at 3 days post treatment (Extended Data Fig. 1a). In order to further optimise treatment conditions for a CRISPRa screen, we analysed the influence of repeated treatment cycles at IC_50_ on cell viability. We found that after three cycles of 100 nM imatinib (day 9), ABCB1 overexpressing cells remained 31.5-fold (sgABCB1-1) and 23.5-fold (sgABCB1-2) more viable than negative control cells (sgNTC) (Extended Data Fig. 1b). These results show that repeated cycles of low imatinib doses allow much greater enrichment of cells with activated resistance genes than a single treatment.

**Figure 1.**
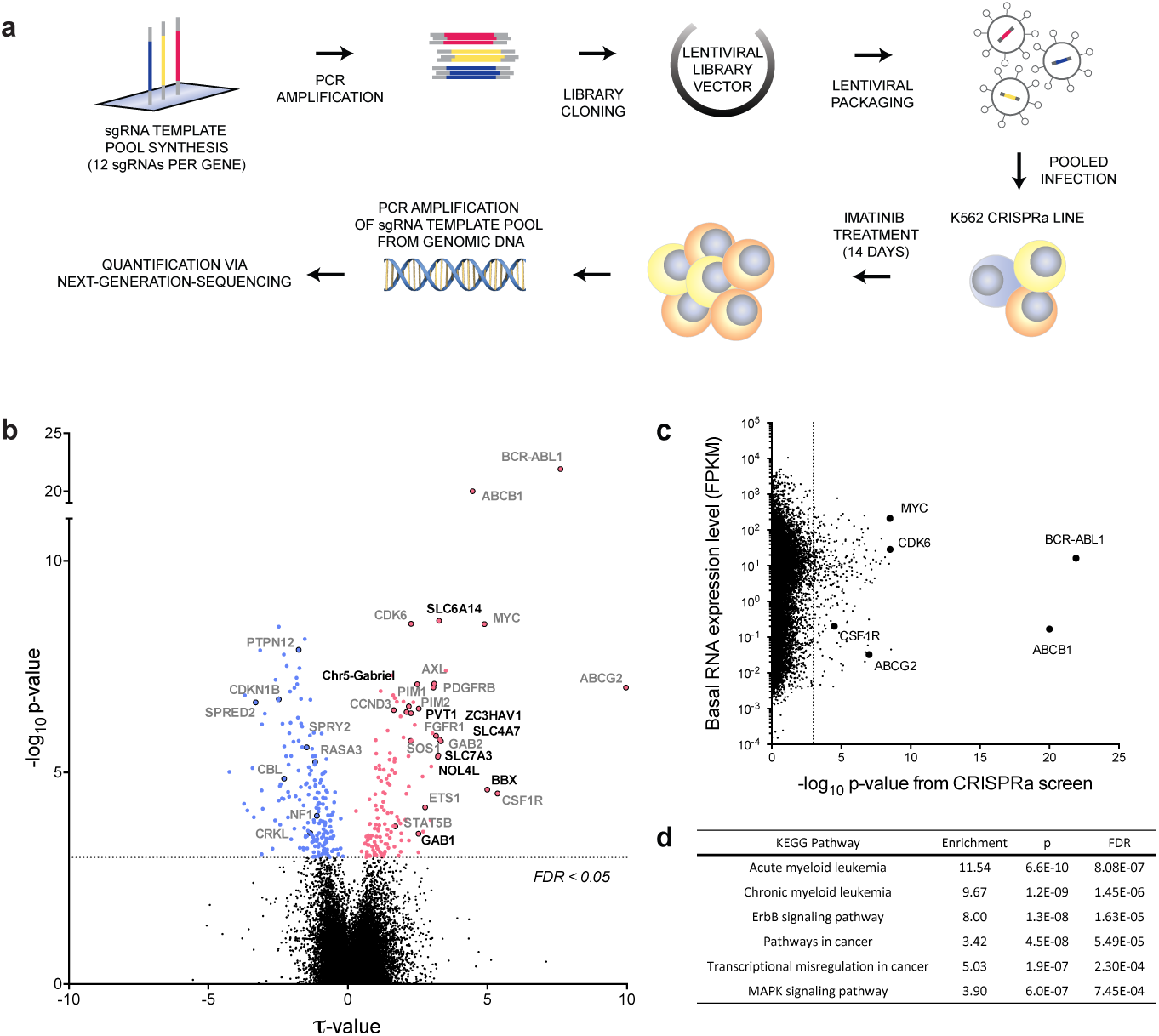
Ultra-complex CRISPRa screen identifies hundreds of genes involved in cancer signalling pathways. **a**, Schematic of genome-scale CRISPRa screening approach (see text for details). **b**, Overview of CRISPRa screen results. Negative τ values indicate a decrease and positive values an increase of cell fitness following gene activation. Significant candidate genes (FDR < 0.05, dotted line) are in colour (blue = decrease, red = increase in fitness). Previously identified gene names are highlighted in gray (see also Fig. 2a) while newly identified and validated candidate genes are labelled in black (see also Fig. 2c). **c,** Comparison of absolute gene expression levels (FPKM) to results from the CRISPRa screen (-logio p-value). **d,** GSEA with the 332 significant candidate genes identified by CRISPRa screen.

To systematically identify genes whose activation can alter the response of cells to BCR-ABL inhibition, we created an ultra-complex, genome-scale sgRNA library consisting of over 260,000 total sgRNAs targeting every Refseq annotated transcript in the human genome, including protein-coding and non-coding transcripts and each isoform thereof individually by up to 12 sgRNAs. The sgRNA library was cloned into a lentiviral sgRNA expression vector and quality controlled via next-generation sequencing. Fidelity of sgRNA sequences was found to be high, with more than 90% perfect alignment rate and narrow distribution of sgRNA sequences, with read counts for 87% of sgRNA sequences falling within one order of magnitude (Extended Data Fig. 2).

**Figure 2.**
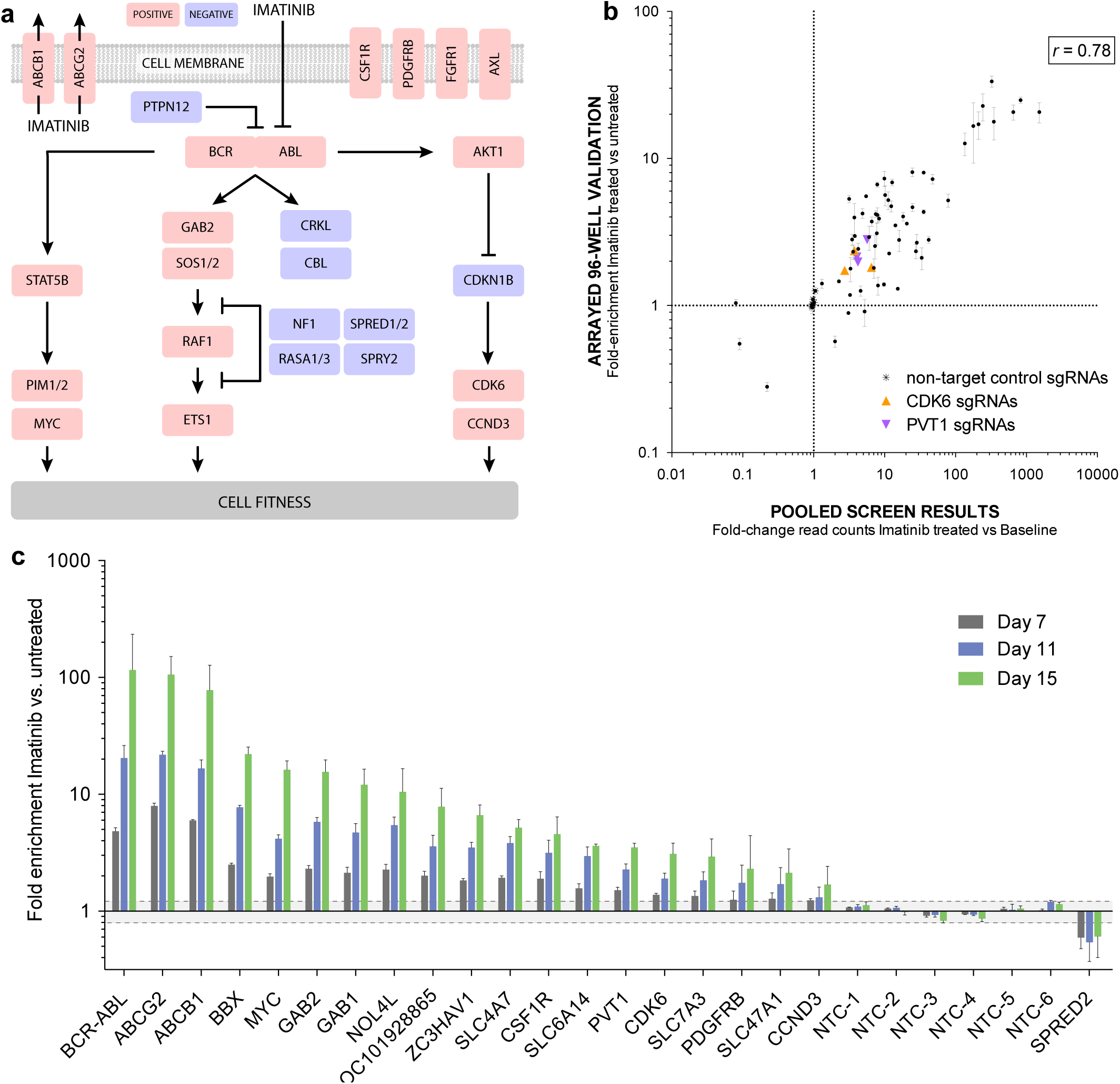
Phenotypes from pooled CRISPRa screen are highly reproducible. **a,** Subset of identified candidate genes mapped onto known signalling pathways (blue = negative, red = positive regulatorof fitness). **b,** sgRNA sequences against 20 significant candidate genes from pooled CRISPRa screen (n=3 sgRNAs/gene) were individually sub-cloned and re-assayed in an arrayed 96-well format. Validation results show high levels of correlation betweeneven subtle phenotypes (e.g. CDK6, PVT1) from the pooled screen. Non-target controls (NTC) show no enrichment. Mean values with s.e.m. are shown. **c,** Candidate gene enrichment was measured over time. Values represent the mean of three sgRNAs targeting each gene with s.e.m. Grey shading = two standard deviations of sgNTCs.

Quality-controlled libraries were packaged into lentiviral particles and used to transduce the K562 CRISPRa target cells at low multiplicity of infection (MOI) as summarised in Figure 1a. The low MOI used for transduction reduced the frequency of multiple-infected cells; hence, one specific gene was activated in each cell. Following antibiotic selection of infected cells, escalating doses of imatinib ranging from IC_50_ to IC_80_ were applied for 14 days to allow cells with altered imatinib tolerance to enrich or disappear from the pool of cells over the course of the screen. Genomic DNA from cells at the beginning and end of the screen was harvested and sgRNA encoding sequences were PCR-amplified from the genomic DNA. Cells that enriched or disenriched, following expression of a certain sgRNA were quantitated via next-generation sequencing (NGS) of the pool of amplified sgRNA template sequences (Extended Table 1). Normalised read count ratios were used by averaging the phenotypes of the top 25% most extreme sgRNAs to compute enrichment scores (τ) and p-values for each gene. Extended Table 2 shows a summary of the results at gene level, alongside the results from an untreated control screen without imatinib as well as the expression levels from each target gene in the K562 CRISPRa cell line used for the screen. We found that out of a total of 26,700 targeted transcripts, the activation of 332 of them significantly (FDR<0.05, p<0.001) altered the fitness of imatinib-treated K562 cells with the overexpression of 57% (188) causing depletion (blue) and 43% (144) enrichment of cells over the course of the screen (Fig. 1b). Activation of 83% of those candidate genes (275) specifically altered the cells’ response to imatinib but had no significant effect (FDR>0.05) on untreated cells while 17% (57) affected cell growth in the imatinib-treated and untreated arm of the screen (Extended Table 2). Phenotypes evoked by the sgRNAs targeting the 332 candidate genes were highly reproducible (r>0.98) between technical screen replicates (Extended Data Fig. 3). Owing to this high reproducibility and the large complexity of the employed sgRNA library (12 sgRNAs/gene), even subtle alterations of cell fitness could be detected at extremely high significance levels; for example, the slight increase in fitness observed after activation of the cell cycle regulators CDK6 (τ = 2.26, p = 3×10^−9^) and CCND3 (τ = 2.18, p = 2.75×10^−7^), or the poorly understood noncoding lncRNA PVT1 (τ = 2.25, p = 4×10^−7^).

**Figure 3.**
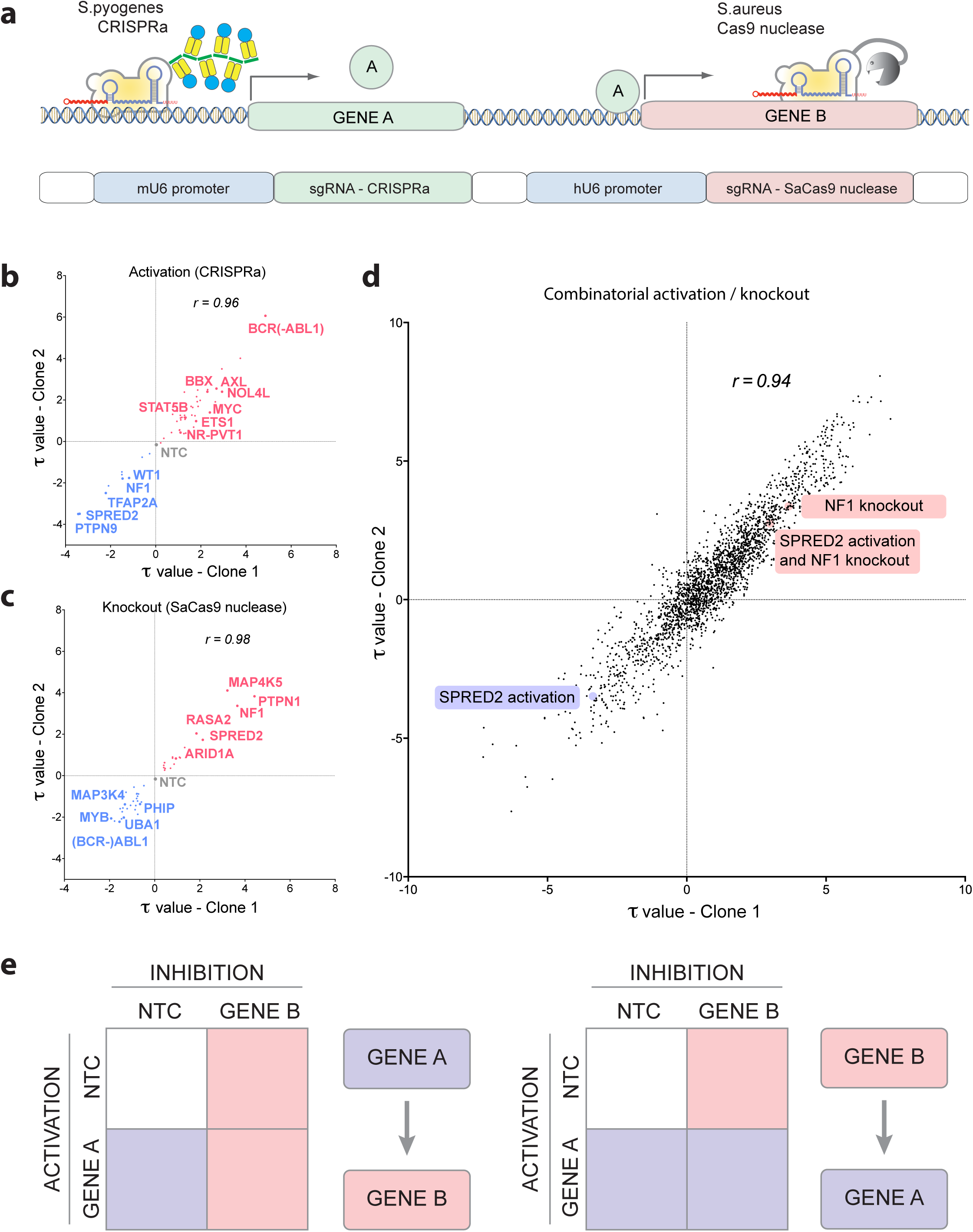
Orthogonal CRISPR/Cas screens can quantify directional genetic interactions. **a,** Schematic of the orthogonal CRISPR/Cas system. CRISPRa system (*S.pyogenes* Cas9-based) was combined with Cas9 nuclease from *S.aureus* and dual sgRNA expression vectors were developed to express appropriate sgRNAs to direct either system to distinct genomic target sites in the same cell. Correlation of τ values from two biological replicates is shown for **b,** gene activation, **c,** gene knockout and **d,** all possible combinations thereof. **e,** if the combinatorial phenotype mimics the knockout phenotype, the activated gene was interpreted to act upstream of the knocked out gene B (left panel), and if the combination mimics the activation phenotype, the activated gene A was interpreted to act downstream (right panel).

An important advantage of the gain-of-function approach used here, as opposed to more commonly employed loss-of-function approaches, is that non-expressed genes can be investigated. To evaluate the relationship between K562 gene expression levels and the ability of CRISPRa to evoke significant fitness phenotypes from targeted genes, we compared FPKM RNA expression levels to gene hit probabilities (Fig. 1c). These analyses showed an approximately five orders of magnitude span in FPKM levels for the top 332 hit genes. These include non-expressed genes (21%, FPKM<10^0^) like ABCB1, ABCG2, or CSF1R which are frequently found activated in drug resistant cancer cells from leukaemia patients^28^, ^29^. Candidate genes also include expressed genes (79% with FPKM>10^0^) with well-studied roles in leukaemia, like MYC, BCR-ABL1 or CDK6^30^. Together these data indicate that imatinib responsive genes could be identified from the full spectrum of gene expression levels, ranging from non-expressed to highly expressed genes.

To assess the quality of the data on a global level, we executed a gene set enrichment analysis (GSEA)^31^, ^32^ of the 332 significantly called genes. GSEA identified the strongest gene enrichment in leukaemia and other cancer-related KEGG signalling pathways (Fig. 1d), supporting the specificity of the CRISPRa screening approach in identifying positive and negative regulators of cancer cell survival pathways. A subset of hit genes from the CRISPRa screens re-assembled the established pathways for Acute Myeloid Leukaemia (AML), reflecting shared core myeloid pathway components with CML. Gene sets related to Erb and MAPK signalling and cancer related pathways also scored significantly, suggesting that the top 332 genes identified in the CRISPRa screen were enriched for key signatures relevant to human cancers. To give a graphic summary of the CRISPRa screen results, candidate genes with known roles in well-established cancer signalling pathways, were assembled (Fig. 2a).

Given the enrichment of key GSEA signatures in the set of identified candidate genes, we tested three sgRNAs against each of the top 20 genes individually in an arrayed 96 well plate validation assay. To quantitate cell survival and proliferation, control cells that expressed no sgRNA were mixed with cells transduced with an individual sgRNA targeting one of the top 20 candidate genes. Because the sgRNA vector carried an mCherry reporter, we conveniently monitored relative cell viability/proliferation over time via a red fluorescence FACS assay, a protocol which we have found to be highly quantitative. The enrichment values from the arrayed validation assay exhibited a high degree of quantitative reproducibility when compared to screen enrichment datafor those sgRNAs (r = 0.78) with a wide dynamic enrichment range over several orders of magnitude (Fig. 2b). Each of the 20 genes showed significant activity compared to the activity of Non-Targeting Control (NTC) sgRNAs (Fig. 2c). Three of the strongest hits, namely ABCB1, ABCG2 and BCR-ABL1 are well known to be overexpressed in CML patients with high tolerance to imatinib^28^. Additionally, we identified BCR-ABL1 binding partners CBL and CRKL^33^, as well as downstream effectors SOS1, SOS2, GAB2, RAF1, MYC, PIM1, PIM2 and STAT5B^34^, the c-Abl phosphatase PTPN12^35^, the Ras-GAPs NF1, RASA1 and RASA3^36^, the cell cycle regulators CDK6^37^ and CCND3^38^ as well as receptor tyrosine kinases with well documented roles in imatinib resistance, such as PDGFRB^39^, FGF1R^40^, CSF1R^29^ and AXL^41^.

### An orthogonal screening system to identify directional genetic interactions

Given the high quality of the CRISPRa screening data and the need to identify functional directional interactions between known and unknown genes, we next established an orthogonal screening system enablingthe simultaneous activation of one gene and inhibition of a second gene (Fig. 3a). Compared to traditional symmetric gene interaction study approaches, the combined gain- and loss-of-function approach results in two opposing outcomes if two genes interact in an activating way. To successfully conduct orthogonal screens, we created a new K562 cell line harbouring the *Streptococcus pyogenes* based CRISPRa system described above, with wild type Cas9 nuclease from *Staphylococcus aureus* (SaCas9). Cas9 proteins from both species have different PAM requirements and Cas9 structural studies found that each enzyme recognises different constant regions of the cognate sgRNA^42^, ^43^. These observations indicate that each Cas9 is not likely to cross-react with the cognate sgRNA engineered for the other orthogonal Cas9 enzyme.

We next wanted to employ the orthogonal CRISPR/Cas system to investigate functional interactions between different genes. For that purpose, we created a combinatorial sgRNA library composed of 2.4 million dual sgRNA vectors. This library consisted of 192 preselected CRISPRa sgRNAs, targeting 87 significantly enriched or disenriched candidate genes from the primary genome-scale CRISPRa screen (2 sgRNAs/gene plus 18 non-target controls) combined with 12,500 SaCas9 nuclease sgRNAs targeting over 1,300 genes for knockout (8 sgRNAs/gene plus 878 non-target controls), including every gene in KEGG annotated cancer-relevant signalling pathways. The orthogonal sgRNA library sequences along with target gene names for the CRISPRa and SaCas9 nuclease system are shown in Extended Tables 3 and 4, respectively.

The dual sgRNA expression library was transduced into two clonal lines of orthogonal K562 cells. The transduced clonal lines were cultured in separate bioreactors in the presence of escalating doses of imatinib. After 19 days, cells were harvested, and sgRNA abundances were analysed from the selected cells via NGS. Individual activation and knockout phenotypes, and all possible combinations thereof were quantified independently from the two clonal screen replicates as previously described^44^. To permit the most rigorous quantification from the determined values, knocked-out genes were only retained for further analysis if the knockout showed a significant phenotype by itself (FDR < 0.05) and if genes were represented by at least two strongly correlating sgRNAs (based on the correlation between genetic interaction patterns, see Methods for details). Likewise, activated genes were retained for further analysis only, if their sgRNA showed correlated genetic interaction patters above a certain threshold (see Methods for details). In this fashion, only the most stringent hit genes were included in the double perturbation analyses.

A necessary value in gene interaction studies is the measurement of single-gene perturbations. To facilitate dissection of activation-only phenotypes and knockout-only phenotypes, we included a large number of non-target control sequences in the combination library (see above). Hence the library contained dual sgRNA vectors which harboured only an activating sgRNA or only an inactivating sgRNA at one position and a non-targeting control sgRNA in the other. Dissecting these two populations of vectors allowed the clean evaluation of replicate performance for both single activation (r=0.96, Fig. 3b) and knockout phenotypes (r=0.98, Fig. 3c). Both single gene activation and single knockout phenotypes included negative and positive regulators of cell fitness in the presence of imatinib and these results were in strong agreement between both replicates (Fig. 3b and c) and the initial CRISPRa screen (Extended Data Fig. 4). Most importantly, we also observed highly reproducible (r=0.94) and quantitative measurements of phenotypes derived from the dual sgRNA vectors combining all possible combinations of sgRNAs (Fig. 3d). All single and combinatorial τ values were clustered hierarchically based on the un-centred Pearson correlations of their τ profile and a map of the results is shown in Extended Data Fig. 5. Together these data confirm: 1) the ability for both Cas systems to work in parallel to produce activation and knock-out phenotypes in the same celland 2) the suitability of our NGS analysis pipeline to accurately quantify phenotypes from combinatorial gene perturbations.

**Figure 4.**
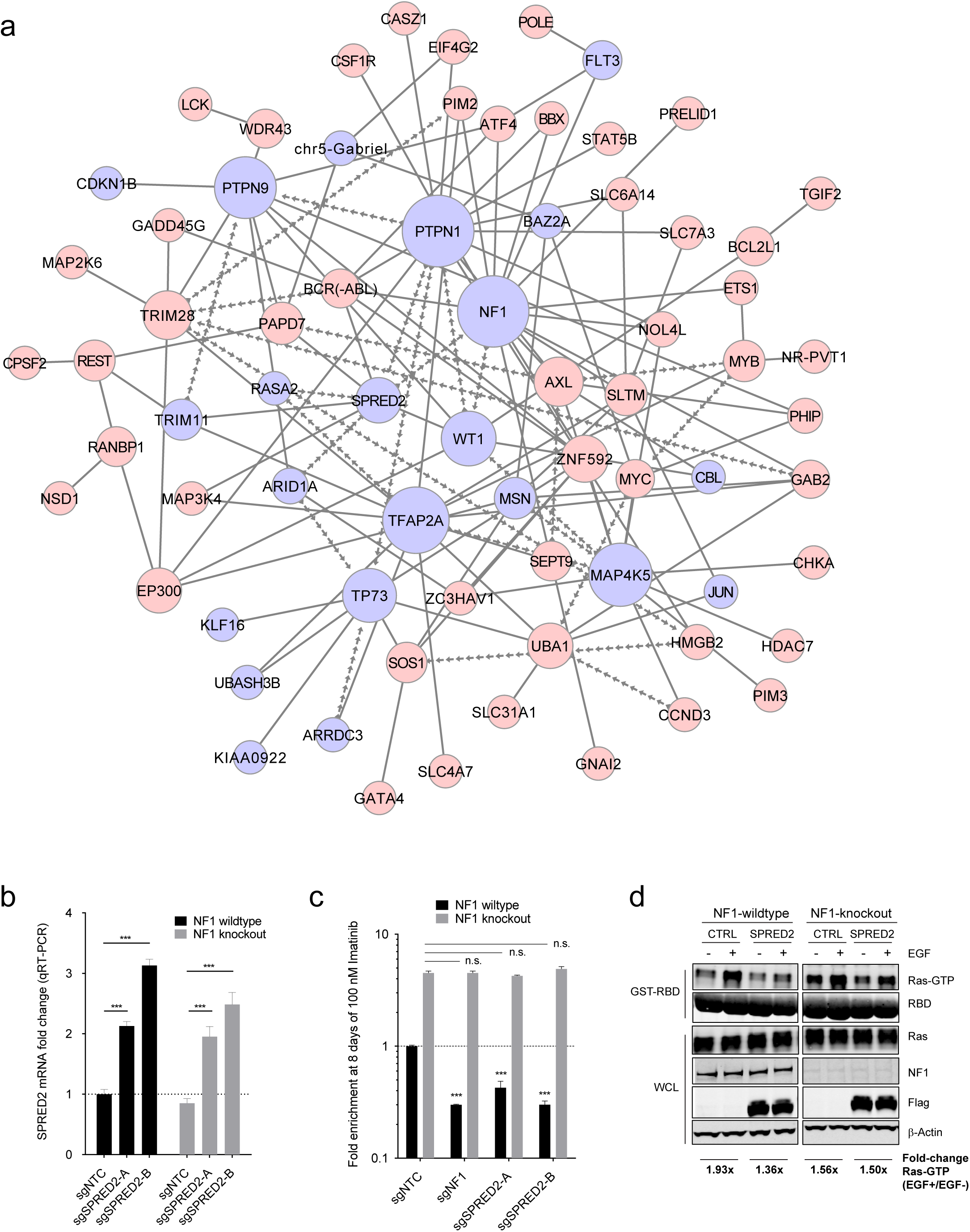
Validation of directional genetic interaction between SPRED2 and NF1. **a,** Based on GI scores determined by the full orthogonal interaction screen, a genetic interaction model was constructed. For positive regulators of cell fitness, nodes are shown in red and for negative regulators in blue. Arrow-shaped edges indicate inferred directional interactions between nodes. Line-shaped edges symbolise genetic interactions where directionality could not be inferred. **b,** Significantly increased SPRED2 mRNA levels were detected via qRT-PCR in NF1 wildtype and knockout cells that expressed SPRED2 activating CRISPRa sgRNAs (mean with s.e.m. is shown, n=3, x-axis indicates expressed CRISPRa sgRNAs). **c,** NF1 and SPRED2 activation significantly sensitises NF1 wildtype but not NF1 knockout cells to imatinib treatment (mean with s.e.m. is shown, n=3, x-axis indicates CRISPRa sgRNAs). **d,** Overexpression of SPRED2 suppresses Ras activation only in NF1-wildtype cells, as judged by Ras-GTP levels. Quantified Ras-GTP level fold-changes of EGF-induced versus non-induced cells are shown. Negative control (CTRL) and SPRED2 overexpression constructs were transiently transfected into HEK293T cells 24h before EGF stimulation.

To systematically identify genetic interactions in the deep dual sgRNA dataset, τ values were used to compute genetic interaction (GI) scores for all gene-gene combinations. For this purpose, all individual gene activation and knockout τ values at the sgRNA level were used to compute expected combinatorial τ values, which were then compared to the measured combinatorial τ values^44^. Genes were considered to interact when their GI scores exceeded a 1x standard deviation cut-off consistently in both biological screen replicates, restricting the resulting dataset to a higher stringency. Directionality of the identified interactions was interpreted, depending on whether the activated or knocked out gene was dominating the combinatorial phenotype (Fig. 3e). To permit quantitative interpretation of directionality in the measured interactions, GI scores and τ values were integrated into a single directionality score Ψ. For genetic interactions between genes having opposing phenotypes, and to maintain stringency, directionality was inferred if Ψ exceeded a 1x standard deviation cut-off. For each of the thousands of interrogated gene-gene combinations, τ values, gene interaction scores and Ψ directionality scores from both clonal replicates are provided in Extended Table 5. Based on the determined τ values and Ψ scores, a directional genetic interaction network that consists of only significant and reproducible interactions determined by our orthogonal screen (Extended Table 6) was constructed and inferred directionality is indicated by arrow heads where possible (Fig. 4a). These data comprise the only reported instance of deduced directionality for a large-scale cancer gene interaction network screen, providing a compendium of directional-edge models. Together, these directional-edge models connect a total of 70 nodes, demonstrating a high level of interconnectivity between nodes. For clarity, interactions from all 70 separated nodes are shown individually in Extended Data Fig. 6.

### SPRED2 sensitises cells through NF1

The orthogonal screen data suggest that the cell growth inhibitive function of SPRED2 is lost with the loss of NF1, suggesting a function of SPRED2 upstream of NF1. NF1 encodes Neurofibromin 1, a Ras-GAP, that switches the Ras protein from its active GTP-bound state into its inactive GDP-bound state, thereby turning off MAPK signalling promoted proliferation^45^. Inactivating NF1 mutations are found at high frequency not only in Neurofibromatosis 1^46^ but also in a variety of cancers like melanoma^47^, ^48^, glioblastoma^49^, ^50^ and ovarian cancer^51^. We first confirmed by qRT-PCR that SPRED2 mRNA levels following CRISPRa mediated gene activation were almost identical in K562 cells with or without NF1 (Fig. 4b). When treated with imatinib, NF1-knockout cells enriched 4.5x, while NF1-wildtype cells with CRISPRa-mediated NF1 activation became 3x more sensitive to the treatment when compared to control cells (Fig. 4c). As expected, the sensitising effect of NF1 activation was reversed in NF1-knockout cells, which showed the same level of enrichment than NF1-knockout control cells without CRISPRa mediated NF1 activation (4.5x). Most importantly, activation of SPRED2 by two different sgRNAs significantly sensitised NF1-wildtype cells to imatinib treatment while in NF1-knockout cells this sensitising effect was fully lost, suggesting that SPRED2 acts exclusively through NF1 in order to sensitize K562 cells to imatinib (Fig. 4c). In support of these results, we show that SPRED2 overexpression suppresses EGF-induced Ras activation in HEK293 cells (1.93x versus 1.36x induction) which is in line with previous findings showing that the double knockdown of SPRED1/2 leads to increased Ras-GTP levels^52^. In addition, we show that the ability of SPRED2 to suppress Ras activation was lost in NF1-knockout cells (1.56x versus 1.5x induction) (Fig. 4d) which further confirms that SPRED2 requires NF1 to exert its sensitising function.

The directional SPRED2-NF1 interaction represents only one example of the genetic interactions identified by our orthogonal Cas9 screen. In addition, a number of genetic interactions observed in our screen have already been described in the literature, including the activation of the MYC gene by the MYB transcription factor^53-55^, the synergistic interaction between MYB and ETS1 in transcriptional regulation of their shared targets^56-58^, dephosphorylation of STAT5B by the phosphatase PTPN1 (also known as PTP1B)^59-61^,and the accepted interaction between PTPN1 and BCR-ABL^62^. Together, these validated and published data confirm the usefulness of this technology to uncover directional genetic interactions between known and unknown factors in cancer cells.

## Discussion

Inferring the direction of genetic interactions in model organisms has been a long-standing challenge. Here, we describe a systematic highly quantitative approach using orthologous CRISPR/Cas enzymes in human cells. While all previous genetic interaction studies have used symmetric systems where the function of both potential interaction partners is lost, the orthogonal approach combines the CRISPR/Cas mediated activation of one interaction partner with the functional loss of a second one. Here, we established the full methodology and reagents necessary to conduct highly parallel dual sgRNA CRISPR/Cas screens, including stable CRISPRa-SaCas9 nuclease cell lines, dual sgRNA libraries and a barcode-free next generation sequencing strategy to quantify sgRNA combinations in orthogonal screens. We demonstrate how this approach can be used to accurately quantify gene activation and knockout phenotypes as well as combinations thereof, to interpret orientation in activating genetic interactions, simply by expressing two sgRNAs, a task that would have been incredibly challenging to achieve before the advent of CRISPR/Cas technology. Moreover, the orthogonal screening approach developed here significantly advances previous approaches that characterise genetic interactions in mammalian cells by means of RNAi^13-15^, ^17^ and complements more recently published symmetric CRISPR/Cas knockout based screening approaches^63^. It also opens the door to combinations of any two CRISPR/Cas-based technologies, such as transcriptional silencing^64^ or targeted DNA methylation^65^, which represents a substantial advance compared to the only other orthogonal CRISPR/Cas-based method published to date by Dahlmann et al., which achieves gene activation and knockout using 'catalytically dead' sgRNAs in combination with a catalytically active CRISPRa Cas9 enzyme from *S.pyogenes^66^*.

Approaches to construct quantifiable directional models have been limited and there have been no established technologies to efficiently specify directionality in pathways. This is particularly a problem in fields such as cancer biology, where a major ongoing focus is to identify synergistic genetic vulnerabilities and directional dependencies that provide a sound basis for the design of rational polytherapies to help prevent drug resistance. We provide a large map of genetic interactions that will help to further understand why some patients respond well to tyrosine kinase inhibitors while others acquire resistances. These dependencies need to be considered when designing a treatment plan for patients harbouring these biomarkers and the described orthogonal platform offers an informed basis by which therapeutic intervention strategies can be designed. Decoding directional genetic interactions through orthogonal CRISPR/Cas screens offers a fresh new approach to uncovering key dependencies in pathways critical for understanding human gene function and disease.

## Acknowledgements

We thank the McManus Lab for helpful support and discussion, Luke A. Gilbert and Marvin E. Tanenbaum for sharing the CRISPRa K562 cell line. This work was supported by NIH awards to MTM (1U01MH105028 and U01CA168370) and ML (GM119139) and the Chan-Zuckerberg Biohub.

---

**Extended Data Figure 1 | a**, Dose response curve on K562-CRISPRa cells with non-target control sgRNA (sgNTC) or sgRNAs targeting the imatinib efflux transporter ABCB1 for activation (sgABCB1-1 and sgABCB1-2). IC_50_ values are shown in brackets. **b,** K562-CRISPRa cells expressing sgRNAs from **a** or no sgRNA treated repeatedly with 100 nM imatinib on days 0, 3 and 6. Fractions of viable cells, SEMs and resistance factors (RF = fraction of viable ABCB1 overexpressing cells / fraction of viable control cells) are shown from days 3, 6 and 9.

**Extended Data Figure 2 |** CRISPRa sgRNA library distribution. Shown is the cumulative fraction of sgRNAs in the CRISPRa library ranked by their normalised read counts. Shaded area highlights one order of magnitude around median read count.

**Extended Data Figure 3 |** CRISPRa screen reproducibility. Correlation between sgRNA phenotypes from three technical screen replicates (R1-3). Shown are fold-changes from sgRNAs (cutoff>100 reads per sgRNA in baseline sample) targeting all significantly dis-/enriched CRISPRa screen candidate genes.

**Extended Data Figure 4 |** CRISPRa τ values from the primary screen and CRISPRa τ values from the orthogonal screen (non-target control in SaCas9 nuclease position) show a high level of correlation (r = 0.9267).

**Extended Data Figure 5 |** Gene activation phenotypes are shown in rows, knockout phenotypes in columns. Colours represent a decrease (blue) or increase (red) in cell fitness following indicated genetic modifications.

**Extended Data Figure 6 |** All genetic interactions identified from the orthogonal CRISPR screen separated by nodes (highlighted in yellow).

## Methods

### CRISPRa and orthogonal K562 cell lines

K562 CRISPRa cells^67^,^68^ were kindly provided by Luke Gilbert and cultured in RPMI 1640 medium, supplemented with 10% fetal bovine serum and 1x Anti-Anti (Gibco). Via lentiviral transduction, *S. aureus* Cas9 under the control of an EF1a promoter, was introduced into K562 CRISPRa cells. Successfully transduced cells were selected with hygromycin (200 ug/mL) and single clones were expanded for 14 days.

### Cell viability assays

Cells were seeded at 10,000 cells per 96-well in 200 uL RPMI-1640 (10% FBS, 1% pen-strep) with indicated imatinib concentrations. Viability was determined at indicated time points by mixing 100 uL cell suspension with 50 uLresazurine medium (50 ug/mL, Acros Organics). After 2h incubation, fluorescence was quantified on a plate reader (BMG Labtech) at excitation: 530 nm and emmision: 590 nm.

### Vector maps

For the single sgRNA (sgLenti), dual sgRNA (sgLenti-orthogonal) and SaCas9 nuclease vector, vector maps are provided in Genbank format (Extended Data 1-3).

### CRISPRa and orthogonal sgRNA library design

For the initial CRISPRa screen, a genome-wide sgRNA library of 260,000 sgRNAs was generated targeting the promoter regions of coding transcripts and selected non-coding regions, including 7,700 non-target control sequences (NTC). The promoter regions for coding transcripts targeted windows 25 to 500bp upstream of the Refseq-annotated transcription start sites. SgRNAs were designed against targets in the promoters that are of the format (N)_20_NGG, and selected sgRNAs must pass the following off-targeting criteria: 1) the 11bp-seed must not have an exact match in any other promoter region, and 2) if there is an exact off-target seed match, then the rest of the sgRNA must have at least 7 mismatches with the potential off-target site. After all sgRNAs that pass off-targeting criteria were generated, up to 12 sgRNAs/transcript were selected that were nearest to the transcription start sites. All sgRNA sequences are shown in Extended Table 1. In addition to the sgRNA sequence, every plasmid contained a unique 20 nt barcode sequence (see sgLenti vector map, Extended Data 1). This sequence allowed the distinction between sgRNAs expressed from different plasmids and hence in different sub-populations of cells and was used to bin cells into mutually exclusive barcode bins to create technical screen replicates after sequencing.

For the secondary genetic interaction screen, a focused nuclease-active *S. aureus* Cas9 library was generated targeting 1327 genes. For the selected genes, sgRNAs targeting coding exons and microRNA hairpins were generated using Cas-Designer ^69^, generating sgRNAs that were adjacent to the PAM sequence ‘NHGRRT’ (H = A, C, or T), which allows for targeting with *S. aureus* Cas9 but not with *S. pyogenes* Cas9. Potential off-targets against the human genome were identified using Cas-OFFinder^66^. Cas-OFFinder sgRNA results were ranked, penalizing sgRNAs that have perfect-seed off-targets and 5 mismatches or less in potential off-target regions. The 20% of sgRNAs with the highest off-target penalties and bottom 20% of sgRNAs with the lowest out-of-frame scores from Cas-Designer were eliminated. From the resulting list of sgRNAs, up to 8 sgRNAs/gene were selected, targeting the most 5 constitutive exons for each gene.

### CRISPRa and orthogonal sgRNA library cloning

The designed 20 nt target specific sgRNA sequences were synthesised as a pool, on microarray surfaces (CustomArray, Inc.), flanked by overhangs compatible with Gibson Assembly^58^ into the pSico based sgLenti sgRNA library vector after AarI (Thermo-Fischer) restriction digest. The synthesised sgRNA template sequences were of the format: 5’-GGAGAACCACCTTGTTGG-(N)_20_-GTTTAAGAGCTATGCTGGAAAC-3’. Template pools were PCR amplified using Phusion Flash High-Fidelity PCR Master Mix (ThermoFisher Scientific) according to the manufacturers protocol with 1 ng/uL sgRNA template DNA, 1 uM forward primer (5’-GGAGAACCACCTTGTTGG-3’), 1 uM reverse primer (5’-GTTTCCAGCATAGCTCTTAAAC-3’) and the following cycle numbers: 1× (98C for 3 min), 15× (98C for 1 sec, 55C for 15 sec, 72C for 20 sec) and 1× (72C for 5 min). PCR products were purified using Minelute columns (Qiagen). Library vector sgLenti was preapred by restriction digest with AarI (Thermo-Fischer) at 37C overnight, followed by agarose gel excision of the digested band and purification using NucleoSpin columns (Macherey-Nagel). Using Gibson Assmbly Master Mix (NEB), 1000 ng digested sgLenti and 100 ng amplified sgRNA library insert were assembled in a total 200 uL reaction volume for 15 min at 50°C. The reaction was purified using P-30 buffer exchange columns (Biorad) that were equilibrated 5x with H_2_O and the total eluted volume was transformed into three vials of Electromax DH5a-E (Invitrogen) using pre-chilled 1 mm cuvettes and 2.0 kV, 200 Ohm, 25 uF on the Gene PulserXcell system (Biorad). Transformed E.coli were recovered, cultured overnight in 500 mL LB (100 ug/mL ampicillin) and used for Maxiprep (Qiagen). In parallel, a fraction of the transformation reaction was plated and used to determine the total number of transformed clones. The coverage was determined to be 70x clones per sgRNA ensuring even representation of all library sgRNA sequences and their narrow distribution (Extended Data Fig. 2).

For orthogonal CRISPR libraries, CRISPRa sgRNA pools were cloned into position 1 of the AarI-digested plasmid sgLenti-orthogonal exactly as described for the CRISPRa library. Following amplification in E.coli, library plasmids with the first position cloned were digested with BfuAI (NEB) to allow cloning of SaCas9 sgRNAs into the second position. To remove undigested orthogonal sgRNA library plasmid from the pool, the purified (Nucleospin, Macherey-Nagel) BfuAI digested plasmid was subsequently digested with AscI for which restriction sites exist in the stuffer sequences in sgRNA positions 1 and 2. BfuAI/AscI digested plasmid was extracted from 1% Agarose gel (Nucleospin, Macherey-Nagel). Synthesised SaCas9 sgRNA template sequences were of the format: 5’-GAAAGGACGAAACACCGTG-(N)_22_-GTTTTAGTACTCTGGAAACAGAATCT-3’. PCR amplification of the SaCas9 template pool was performed as described above using primer sequences: 5’-GAAAGGACGAAACACCGTG-3’ and 5’-AGATTCTGTTTCCAGAGTACTAAAAC-3’ and the purified PCR product was cloned into BfuAI digested sgLenti-orthogonal via Gibson Assembly as described above. The resulting orthogonal sgRNA library was transformed into Electromax cells at 30x coverage as described above and the plasmid sgRNA library pool was purified (Qiagen Plasmid Maxi kit). From the resulting plasmid pool, sgRNA sequences were recovered via PCR as described below and sequenced for quality control. At a read depth of 94x, 2.389 million out of the total possible 2.394 million combinations (>99%) were read at least once, with less than 5% of the library elements read 20 or less times.

### Lentivirus production

HEK293T cells were seeded at 65,000 cells per ccm in 15 cm dishes in 20 mL media (DMEM, 10% fetal bovine serum) and incubated overnight at 37C, 5% CO_2_. The next morning, 8 ug sgRNA library plasmid, 4 ug psPAX2 (Addgene #12260), 4 ug pMD2.G (Addgene #12259) and 40 uLjetPRIME (Polyplus) were mixed into 1 mL serum free OptiMEM (Gibco) with 1x jetPRIME buffer, vortexed and incubated for 10 min at RT and added to the cells. 24 h later, 40U DNAseI (NEB) were added to each plate in order to remove untransfected plasmid and at 72h post-transfection, supernatant was harvested, passed through 0.45 um filters (Millipore, Stericup) and aliquots were stored at −80C.

### Genome-wide and orthogonal CRISPR screens

K562 CRISPRa/orthogonal cells were transduced with lentivirally packaged sgRNA libraries at MOI=0.3 and 500x coverage and cultured in RPMI with 10% FBS and 1x Anti-Anti (Gibco) in a 37°C incubator with 5% CO_2_. 48h post transduction, cells were selected with puromycin (2 ug/mL) for 96h. Following selection, aliquots of 300 million cells each, were frozen down in FBS with 10% DMSO for analysis via NGS (see below). Fully selected cells (300 million) were transferred into a 14 liter CelligenBlu bioreactor (Eppendorf) and sub-cultured at 37°C, pH=7.4 and 2% oxygen. Coverage at cell level was kept above 1000x throughout the whole screen and the culture medium was replaced when cell density reached 1 mio/mL.

For the genome-wide imatinib CRISPRa screen: 14 days post transduction, aliquots of 300 mio cells each were frozen down (baseline sample) as described above and and the an IC_50_ concentration of 100 nM imatinib (Sigma) were added to the bioreactor vessel. Imatinib was refreshed on day 17 (IC_60_ = 150 nM) and day 19 (IC_80_ = 300 nM) and cells for the analysis of the final time point were harvested on day 28. For the untreated control screen, cells were cultured under identical conditions for 14 days but without the addition of imatinib. For the orthogonal genetic interaction screen: Puromycin selected cells (2 ug/mL) at 8 days post transduction (2.5 billion per sample) were frozen down as described above and 100 nM imatinib (IC_50_) were added to the bioreactor vessel. Imatinib concentrations were increased throughout the screen to the IC_60_ concentration of 150 nM (day 10), the IC_80_ of 300 nM (day 13 and 15) and finally the IC_90_ of 500 nM (day 17). On day 19 2.5 billion cells per sample were harvested for downstream analysis via NGS.

### Genomic DNA (gDNA) extraction

Cell pellets from baseline and imatinib treated or end point (untreated) samples were resuspended in 20 mL P1 buffer (Qiagen) with 100 ug/mL RNase A and 0.5% SDS followed by incubation at 37C for 30 min. After that, Proteinase K was added (100 ug/mL final) followed by incubation at 55C for 30 min. After digest, samples were homogenised by passing them three times through a 18G needle followed by three times through a 22G needle. Homogenised samples were mixed with 20 mL Phenol:Chlorophorm:Isoamyl Alcohol (Invitrogen #15593-031), transferred into 50 mL MaXtract tubes (Qiagen) and thoroughly mixed. Samples were then centrifuged at 1,500g for 5 min at room temperature (RT). The aqueous phase was transferred into ultracentrifuge tubes and thoroughly mixed with 2 mL 3M sodium acetate plus 16 mL isopropanol at RT before centrifugation at 15,000g for 15 min. The gDNA pellets were carefully washed with 10 mL 70% ethanol and dried at 37C. Dry pellets were resuspended in H_2_O and gDNA concentration was adjusted to 1 ug/uL. The degree of gDNA shearing was assessed on a 1% agarose gel and gDNA was sheared further by boiling at 95C until average size was between 10-20 kb.

### PCR recovery of sgRNA sequences from gDNA

Multiple PCR reactions were prepared to allow amplification of the total harvested gDNA from a 1000x cell coverage for each sample. For the first round of two nested PCRs, the total volume was 100 uL containing 50 ug sheared gDNA, 0.3 uM forward (5’-ggcttggatttctataacttcgtatagca-3) and reverse (5’-cggggactgtgggcgatgtg-3’) primer, 200 uM each dNTP, 1x Titanium Taq buffer and 1 uL Titanium Taq (Clontech). PCR cycles were: 1x (94C - 3 min), 16x (94C - 30 sec, 65C – 10 sec, 72C – 20 sec), 1x (68C – 2 min). All first round PCRs were pooled. The total volume of the second round PCR was 100 uL containing 2 uL pooled first round PCR, 0.5 uM forward (5’-AATGATACGGCGACCACCGAGATCCACAAAAGGAAACTCACCCTAAC-3’) and reverse (5’-CAAGCAGAAGACGGCATACGAGAT-(N)6-GTGACTGGAGTTCAGACGTG-3’) primer where (N)_6_ is a 6 nt index for sequencing on the Illumina HiSeq platform, 200 uM each dNTP, 1x Titanium Taq buffer and 1 uL Titanium Taq (Clontech). PCR cycles were: 1x (94C - 3 min), 16x (94C - 30 sec, 55C – 10 sec, 72C – 20 sec), 1x (68C – 2 min). The resulting PCR product (344 bp) contained adapter sequences compatible with Illumina Hiseq sequencing platforms and was extracted from a 1% agarose gel. For the orthogonal genetic interaction screen, conditions for the first round PCR were slightly modified to: total reaction volume 80 uL containing 20 ug sheared gDNA and the second round PCR product was 887 bp.

Gel extracted bands from the primary CRISPRa screen were submitted for sequencing on an Illumina HiSeq 2500 platform using paired end 50 kits with the custom sequencing primer 5’-GAGACTATAAGTATCCCTTGGAGAACCACCTTGTTGG-3’ for reading the sgRNA sequence and the Truseq Illumina reverse primer to read out 20 nt random barcode sequences used for generation of technical screen replicates (separation of sgRNA reads into three groups with mutually exclusive barcode sequence bins). For orthogonal dual sgRNA library analysis, single end 50 kits were used and read cycles were split, 25 cycles for Read1 with the sequencing primer above (reading the *S.pyogenes* sgRNA) and 25 read cycles for the Index read with the custom indexing primer 5’-TTGGCTTTATATATCTTGTGGAAAGGACGAAACACCGTG-3’ (reading the *S.aureus* sgRNA).

### Data analysis

Total read counts of sgRNA sequences from each NGS sample were collapsed and quantified via alignment to the sgRNA library reference sequences using Bowtie 2.0^28^. Data analysis was conducted similarly as described previously^14^, ^44^. Briefly, for the primary CRISPRa screen, the frequency of sgRNAs was determined by deep sequencing and the average read count of three technical replicates was used. The phenotype *τ* was calculated to quantify the effect of an sgRNA on cell growth in the presence of imatinib. Specifically, *τ* was calculated as:

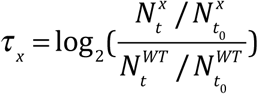
where N^x^ denotes the frequency of sgRNA x and N^WT^ denotes the frequency of non-targeting control sgRNAs before (t_0_) or after (t) imatinib treatment. Gene-level phenotypes were calculated by averaging the phenotypes of the top 25% most extreme sgRNAs targeting this gene. The statistical significance for each gene is determined by comparing the set of *τ* s for sgRNAs targeting it with the set of τ s for non-targeting control sgRNAs using the Mann-Whitney U test, as described previously^44^. To correct for multiple hypothesis testing, we first performed random sampling with replacement among the set of τ s for non-targeting control sgRNAs and calculated P value for each sampling. Then, we calculated the false discover rate (FDR) based on the distribution of P values for all genes in the library and for non-targeting controls generated above. The P-value cutoff was chosen based on an FDR < 0.05.

For the orthogonal double-sgRNA screen, combinations of non-targeting control sgRNAs served as negative control, combinations of one non-targeting control sgRNA and one targeted sgRNA were used to determine single-sgRNA phenotypes and combinations of two targeted sgRNAs were used to calculate double phenotypes. To construct high quality genetic interaction (GI) maps, we implemented a series of filtering steps on the sgRNA level. First of all, on the SaCas9 nuclease side, P values were calculated for each gene as described above. Only the sgRNAs targeting genes that have significant cutting phenotypes (P value <cutoff) were retained. Subsequently, GI scores were calculated using the ‘force-fit’ definition for genetic interactions on the sgRNA level and sgRNAs were further filtered by GI correlation as described previously^44^. On the CRISPRa side, if two sgRNAs targeting the same gene have low correlation, the gene was excluded for further analysis. After the filtering process, gene-level phenotypes and GI scores were calculated by averaging all double-sgRNAs targeting the same gene-gene combinations. Two biological replicates were analysed separately and were combined for the averaged τ map and GI map. Genes were clustered hierarchically based on the uncentered Pearson correlations of their τ profile using Cluster^70^ and visualised by TreeView^71^.

### Directional genetic interaction network model

Genetic interactions whose GI scores exceeded a 1x standard deviation consistently in both clonal screen replicates were used to construct a GI network (Extended Table 6). To quantify directionality in genetic interactions, a directionality score (Ψ) was calculated as Ψ = (τ_activation_)x(τ_knockout_)x(GI)^2^, resulting in a negative Ψ when gene activation and knockout have opposing phenotypes. Negative Ys below a 1x standard deviation of all calculated Ys were used to infer the direction of genetic interactions. The network analysis software platform Cytoscape^72^ was used to visualise the genetic interaction model. Where applicable, directionality in GIs was indicated by arrow shaped edges and line shaped edges indicate significant GIs for which directionality could not be inferred. Nodes were coloured according to gene function with blue symbolizing genes that act to decrease and red to increase cell fitness.

### Arrayed competitive growth validation experiments

Individual CRISPRa or orthogonal dual sgRNA sequences for validation experiments were sub-cloned into the same vector as the respective libraries using the library cloning protocols described above. All library vectors coexpressed mCherry which was used to track the abundance of sgRNA expressing cell populations in growth competition assays. For this purpose, sgRNA expressing cells were mixed with parental - mCherry-negative - cells at ratios between 1:1 and 1:3 in 96-well plates before repeated treatment with 100 nM imatinib for up to 15 days. Enrichment or depletion of the mCherry positive (sgRNA expressing) cell population, indicating an increase or decrease of imatinib resistance following sgRNA expression, could conveniently be followed via FACS quantification of the mCherry-positive versus mCherry negative population. Enrichment factors were calculated as follows: (mCherry-positive/mCherry-negative)IMATINIB/(mCherry-positive/mCherry-negative)_UNTREATED_. Each value was quantified from three separate 96-wells.

### Quantitative RT-PCR

Total RNA from sgRNA expressing cells was purified using Rneasy Mini columns (Qiagen). Taqman probe assays (Applied Biosystems) were used with FAM labelled probes for target genes and VIC labelled probes for the housekeeping gene HPRT1. Reactions were carried out using the one step qRT-PCR master mix TaqMan RNA-to-C_T_ (Applied Biosystems) according to the manufacturer’s instructions on the 2900 HT Fast RT-PCR machine (Applied Biosystems).

### Western blot analyses

HEK 293T cells were transfected with indicated flag tagged plasmids, serum starved for 24 hours, and stimulated with EGF. Ras-GTP was assessed by GST-Raf1 RBD. HEK 293T cells were transfected with pcDNA3.1 Flag-eGFP (CTRL) and Flag-SPRED2 (SPRED2) using Lipofectamine 2000 (ThermoFisher Scientific, 11668019), serum starved for 24 hours, and stimulated with 20ng/ml recombinant human EGF (Invitrogen, PHG0311). Cells were washed with PBS and lysed in TNM buffer (0.2 M Tris pH 7.5, 1% Triton X-100, 1.5M NaCl, 50 mM MgCl2, 1mM DTT, protease and phosphatase inhibitor cocktails. Lysate was cleared and 1,000ug protein was subject GST-Raf1 RBD agarose beads (McCormick lab, in house) for 1.5 hours. Samples were analysed by Western blot using the following antibodies: *NF1* (SCBT, sc-67 [D]), Flag (Sigma, F1804), pan-Ras (Cytoskeleton, Inc, AESA02), β-Actin (Sigma, A5441). *NF1*-Null HEK 293T cells were generated using Cas9 and sgRNA targeting exon 2 with the sequence AGTCAGTACTGAGCACAACA (Shalem, O., et al., 2013). Following single cell cloning, target sequence amplification by PCR, TOPO cloning, and Sanger sequencing, both NF-1 alleles were confirmed deleted by a 1bp insertion resulting in *NF1* (N39fs) and a 11bp deletion resulting in *NF1* (S35fs).

